# Building reusable phage and antibiotic treatments via exploitation of bacteria-phage coevolutionary dynamics

**DOI:** 10.1101/2021.03.31.437900

**Authors:** James Gurney, Sam P. Brown

**Author notes:** Correspondence &.

## Abstract

People with chronic (long-lasting) infections face the problem that treatment options diminish in time as the pathogen evolves increasing resistance. To address this challenge, we exploit phage and bacterial co-evolution, producing dynamic selection pressures that can return the pathogen to a state of susceptibility to the initial (regulator-approved) therapy. We show that phage OMKO1 alone triggers Arms Race Dynamic (ARD) co-evolution with the pathogen *Pseudomonas aeruginosa*, leading to generalized phage resistance and crucially – failure at reuse. In contrast, co-administration of the phage with antibiotics triggers Fluctuating Selection Dynamics (FSD) co-evolution, allowing for effective reuse after 20 days of treatment. We pursue medical relevance in our experiments with the use of clinically important pathogens, antibiotics, phage, and a benchmarked synthetic sputum medium. Phenotypic and genomic characterization of evolved isolates demonstrates that efflux-targeting phage OMKO1 exerts continued selection for antibiotic susceptibility regardless of co-evolutionary dynamic or antibiotic co-treatment, opening the door for evolutionary robust phage therapy.

## Introduction

Bacterial infections in otherwise healthy people are usually rapidly resolved by effective immune responses, potentially aided by antibiotics [1]. In some cases, however, infections fail to resolve even with treatment, leading to the establishment of a chronic infection, imposing elevated morbidity and mortality on affected individuals[2]. Chronic infections are a rising burden on health-care systems due to increasing populations of people at-risk, including the elderly, and people with co-morbidities such as diabetes and heart disease[3]. Management of chronic infections is further complicated by the capacity of pathogen lineages to evolve during the course of infection [4]. In the context of failing antibiotics, there is renewed interest in alternative treatments for chronic infections, including bacteriophage (phage) therapy [5, 6].

Phage are viruses that infect bacteria and are highly abundant in all environments [7-12]. Rising antibiotic resistance levels has led to a greater investigation of novel treatment alternatives, including phage therapy [13-16]. Phage have potential to act as bactericidal treatments alone, and in combination with conventional therapies. However, phage therapy has limitations. Most phage have narrow host ranges (narrow spectrum, in antibiotic terms), which can be overcome by the use of phage cocktails, expanding the host range but also increasing the complexity of regulatory approval by the FDA or other bodies [17]. A second challenge is that the evolution of bacterial resistance to phage attack is rapid, reducing the degree of bacterial lysis and therefore immediate treatment efficacy [13, 18]. Even in the context of cocktail treatments this rapid and predictable loss of efficacy is of particular concern in chronic infections due to the longer timescale of infection control [19, 20].

Phage as a therapeutic are different from antibiotics as phage have the potential to evolve countermoves to bacterial resistance: they can co-evolve with bacteria [18, 21-23]. It is possible that the first bacterial move is a ‘check mate’, for instance loss of surface receptor leading to phage extinction [24]. Yet in some cases, phage countermoves are possible leading to ongoing co-evolutionary dynamics and ongoing killing and evolutionary steering of the pathogen by the phage therapeutic [25]. Here we propose that the specific nature of co-evolutionary dynamics has important consequences for both the efficacy of phage treatment and the potential re-usability of a defined (and therefore more easily FDA-approvable) phage therapy.

In light of established evolutionary theory [18, 26], there are two co-evolutionary dynamics of importance for this work – escalating ‘Arms Race Dynamics’ (ARD) and diversifying ‘Fluctuating Selection Dynamics’ (FSD) [27]. Both have been shown to occur in phage-bacteria interactions [22, 23, 28, 29]. The ARD model captures directional, progressive arms-race evolution where phage get better at infecting ancestral bacteria while bacteria get better at resisting ancestral phage. The outcome of ARD is comparable to selection for resistance against antibiotics; through time bacteria evolve increased resistance to a defined treatment. Under antibiotic selection bacteria develop resistance to the antibiotic, it is a static target for evolution to move towards. Under phage selection the bacteria also develop resistance to the ancestor phage. This increase in resistance therefore reduces the ability of repeated application. In terms of phage therapy this would limit the reuse of a licensed phage therapeutic, raising particular concerns in the context of chronic infections. In contrast, FSD captures a diversifying evolutionary process where both bacteria and phage cycle through many modes of interaction with phage profiting and bacteria suffering from specific ‘lock and key’ fits. Importantly, if phage and bacteria enter into an FSD cycle then ancestral phage (the licensed phage) may be reusable.

A growing number of experimental evolution studies have demonstrated that co-evolutionary dynamics can transition from arms-race dynamics into fluctuating selection dynamics in nutrient limited environments [18, 30]. For example, while the bacterium *Pseudomonas fluorescens* and the phage SBW25φ2 show arms-race co-evolution in rich lab media (King’s B media), in a simple soil supplemented M9 media the dynamics transition to FSD [23]. A more nutrient limited environment could in principle curtail ARD through mutation limitation, yet a follow-on study reported that bacterial isolates with generalized resistance (resistance to phage from multiple time points) emerged but remained at low frequencies [30]. Together these results indicate that the transition from ARD to FSD is governed by context-dependent costs that limit the emergence of generalized resistance and ARD.

In this study we place these foundational evolutionary results in the applied context of developing novel phage therapy strategies for chronic infections. If phage and their bacterial targets experience arms-race dynamics, then re-administration of a treatment will become less effective over time due to the emergence of generalized resistance. However, under FSD the temporal ‘lock and key’ process allows in theory for effective re-administration of an ancestral (potentially FDA approved or patentable) phage if required. The question then becomes an evolutionary one – how can we encourage fluctuating selection dynamics? We hypothesize that co-administration of phage with antibiotics will promote FSD co-evolution – and consequently an improved efficacy on secondary use of the ancestral phage preparation.

To test this hypothesis in a relevant chronic infection context, we focus on a model chronic infection system: the notorious human pathogen *Pseudomonas aeruginosa* (PA) in cystic fibrosis (CF) lung infections. CF is a genetic disorder that leads to lifelong chronic lung infections, with PA the leading cause of mortality [31]. In chronic infections such as CF, the ability to reuse treatments is vitally important. Using a synthetic sputum media (SCFM2) which captures both the key nutrients of CF sputum and the *in vivo* transcriptional profile of PA [32-34]. We co-evolved the phage OMKO1 and PA for 20 days in the presence or absence of 3 clinical antibiotics (ciprofloxacin (Cip), tobramycin (Tob), and piperacillin/tazobactam (Pip)) at concentrations found in sputum during treatment in CF patients. We find that co-evolutionary dynamics shift from ARD to FSD when the drugs are present, and the shift to FSD is associated with substantial improvements on secondary use efficacy, particularly when phage is co-administered with ciprofloxacin. The specific phage we use (OMKO1) has been demonstrated to impose selection against antibiotic resistance by targeting bacterial efflux [6, 35] a strategy we have recently labeled ‘phage steering’ [25, 36]. We further demonstrate that the anti-efflux (and antibiotic sensitizing) impact of OMKO1 continues in this multi-component treatment context and works in concert with the reuse advantages of triggering FSD co-evolution.

## Results

### Co-evolutionary dynamics shift in response to antibiotics

To begin our analysis, we first ask – what are the co-evolutionary dynamics of phage OMKO1 and reference PA strain PAO1 in a synthetic sputum environment (SCFM2) that mimics the physiological conditions of sputum [33, 34], and in the absence of antibiotics. Previous experimental work demonstrates that co-evolutionary dynamics can be sensitive to specific nutrient environments, leaving open the possibility that the idealized ARD will not be present in this specific environment.

To test for the presence of ARD, we conducted 20 days of experimental evolution, taking frozen samples of bacteria and phage every 2 days (10 passages; T1-T10) in order to allow subsequent ‘time-shift’ experiments [18] to assess phage infectivity and bacterial resistance across isolates taken through evolutionary time (passages T1 ,T5, T9). Using the time-shift approach, we found a clear signature of ARD (Figure 2), illustrated by an increasing bacterial resistance through evolutionary time to a constant phage reference (Figure 2A) and an increasing phage infectivity to a constant bacterial reference (Figure 2B). To formally test for ARD, we use a generalized linear mixed effect method (GLMM) with phage time and bacterial time as fixed effects and the replicate lines as random effects [38]. This model identifies an escalating arms race, through the significant positive co-efficient for the time shift (see SI table 1). In other words, the phage got better at infecting through time and the bacteria got better at resisting through time. The Figure 2A result highlights the applied challenge in the clinical reuse context: bacterial co-evolution displays a clear trajectory towards generalized resistance, increasingly effective against earlier phage isolates.

**Figure 1.**
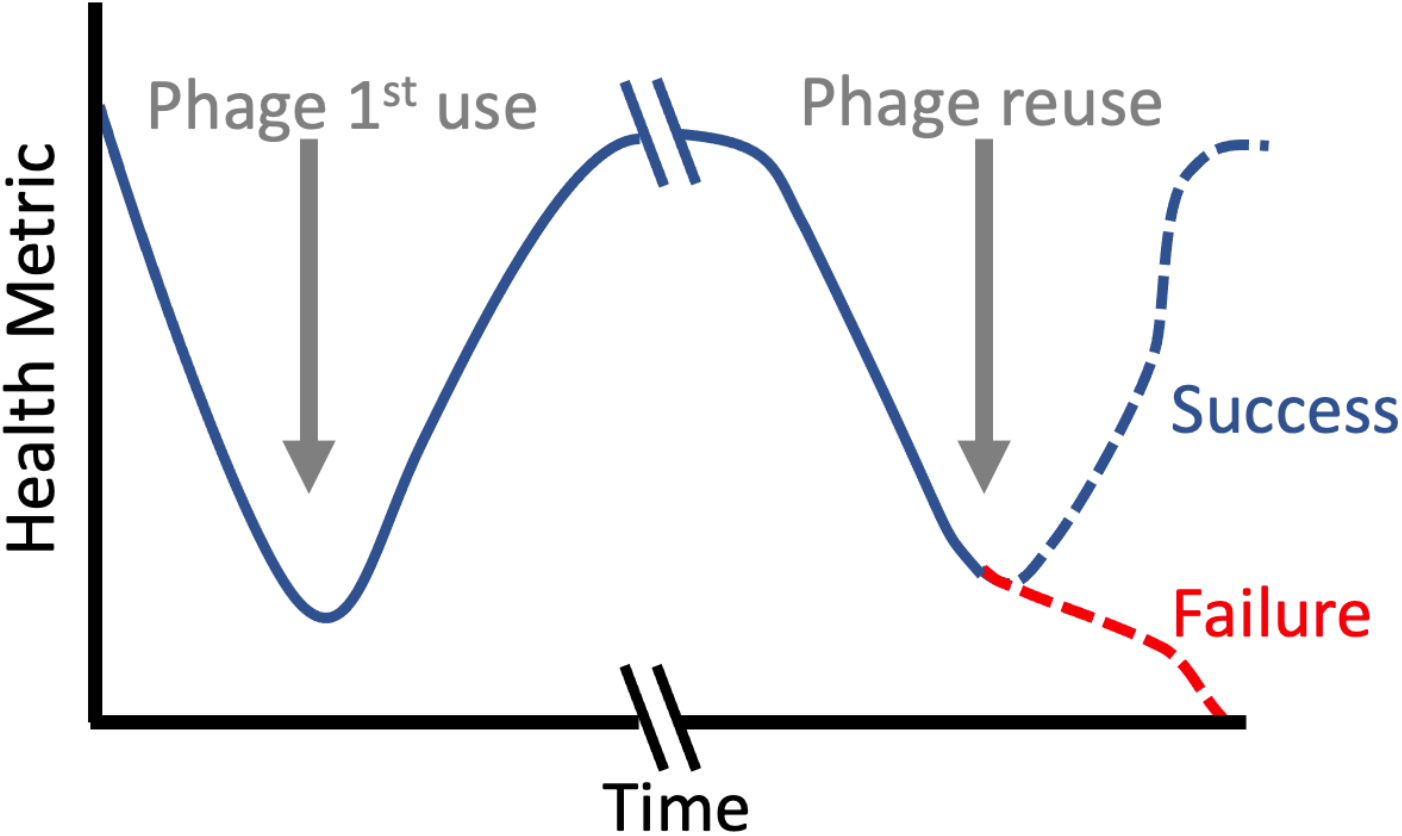
Schematic of phage therapy and potential for reuse. Introduction of phage therapy (grey arrow) reduces bacterial density leading to an improved patient health metric (e.g. lung function in the context of chronic Cystic Fibrosis treatment [37]). Chronic infections typically require repeated treatment, so we ask: is the initial phage preparation reusable? Under an arms-race coevolutionary dynamic, we expect general resistance to fix following the initial round of therapy. Reintroducing the original phage would then impose minimal control on the pathogen, leading to a state of clinical failure (red dashed line). In contrast, a fluctuating selection dynamic in response to the initial treatment could result in an evolved pathogen that is then susceptible to the ancestral and licensed phage, resulting in a repeated clinical success (blue dashed line).

**Figure 2.**
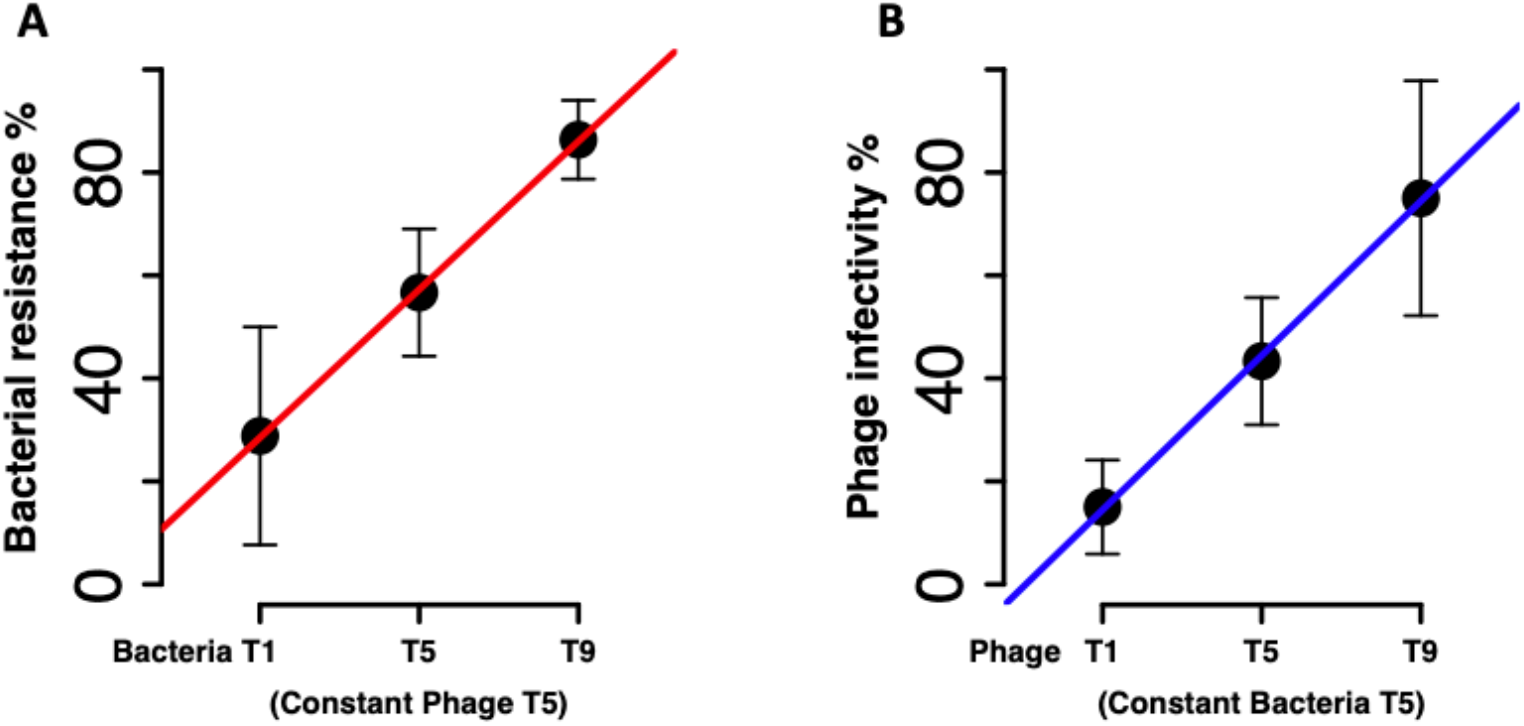
Arms Race co-evolutionary dynamics in a clinically relevant environment SCFM2. Using our frozen ‘fossil record’ of bacteria-phage co-evolution we conducted ‘time-shift’ experiments [18] to assess the performance of bacteria and phage against their co-evolutionary opponent from the past, present or future. We track time in passage transfers, T (transfers were conducted every 2 days). (A) Bacterial % resistance through evolutionary time against a constant phage (T5). (B) Phage % infectivity through evolutionary time against a constant (T5) bacterial passage. Both bacteria (A) and phage (B) show progressive improvements through evolutionary time, versus a time-constant challenger, supporting the Arms-Race Dynamic (ARD) model. This implies a growing generalized resistance that will preclude reuse of an ancestral phage. Error bars are 95% Confidence intervals.

We next ask how the addition of antibiotics (in addition to the phage OMKO1) modifies the co-evolutionary dynamic of bacteria and phage. Antibiotics were reapplied each passage mimicking repeated clinical administration. Figure 3 follows the same format as Figure 2, however the data illustrates a clear absence of progressive, arms-race dynamic co-evolution under all antibiotic treatments. Our statistical (GLMM) analysis rejects an ARD model due to the absence of progressive improvements in bacterial resistance or phage infectivity. In contrast we find support for the FSD model which predicts infectivity to be the result of significant interactions between phage time and bacterial time for all antibiotic co-treatments (SI table 2). [30].

**Figure 3.**
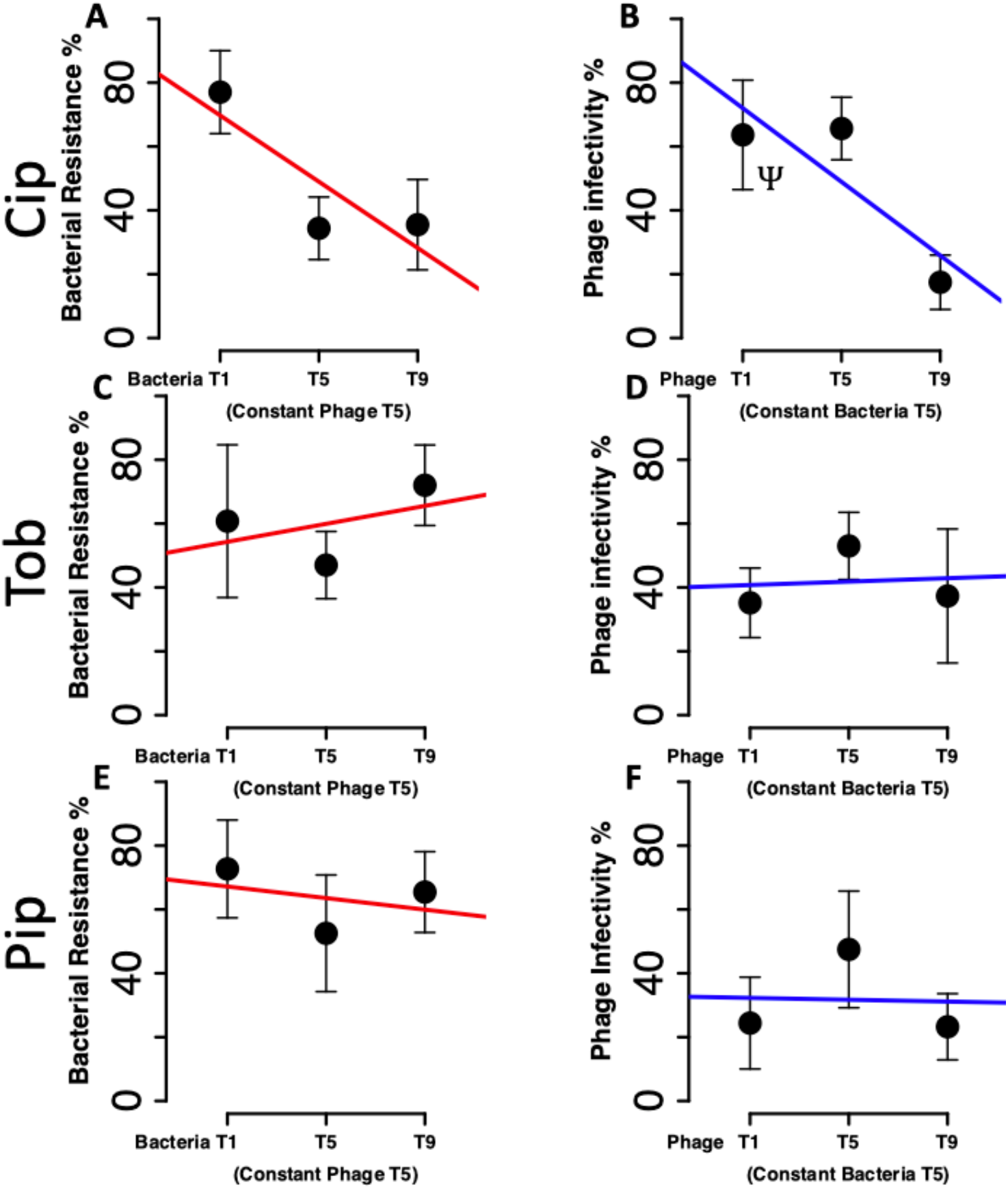
Co-evolutionary dynamics shift in response to antibiotics. Using the same time shift experimental approach as in Figure 2, we now assess the impact of antibiotic co-administration. (A,B) Addition of ciprofloxacin and phage. (C,D) Addition of tobramycin and phage. (E,F) Addition of piperacillin/tazobactam and phage. In the presence of antibiotic co-administration, we do not find progressive improvements through evolutionary time, versus a time-constant challenger. Error bars are 95% Confidence Interval.

### Phage combined with antibiotics promotes phage reuse

Figures 2, 3 indicate that antibiotic co-administration will shift co-evolutionary dynamics of PAO1 and OMKO1 in SCFM2 from ARD (Figure 2) to FSD (Figure 3). In a clinical re-use context, Figure 2 highlights the difficulty of ARD: the T9 (evolved) bacteria shows high (>80%) resistance to even a recent (T5) phage; and conversely an earlier (T1) phage shows poor efficacy (<20%) on an intermediate (T5) bacterium. Figure 3 shows that these values are improved given antibiotic co-administration. In particular, an early (T1) phage isolate under ciprofloxacin coadministration showed over 60% infectivity against an intermate (T5) bacterial isolate (Figure 3B), suggestive of the potential for ancestral phage reuse. We now directly test this prediction by comparing the level of resistance to the initial ancestral (T0) phage in evolved bacterial isolates from the final T10 passage (Figure 4). When evolved under phage alone, we observed high resistance to the ancestor phage, consistent with the ARD dynamics (Figure 2). With phage plus antibiotics, there is partial susceptibility to the ancestral phage, consistent with the FSD seen in figure 3. The ciprofloxacin + OMKO1 line had the largest effect, with ancestral phage infectivity approaching the level seen on the ancestral bacteria (Pink region in Figure 4). This suggests that co-administration of OMKO1 with ciprofloxacin holds the potential to overcome the ARD-barrier to phage therapy reuse.

**Figure 4.**
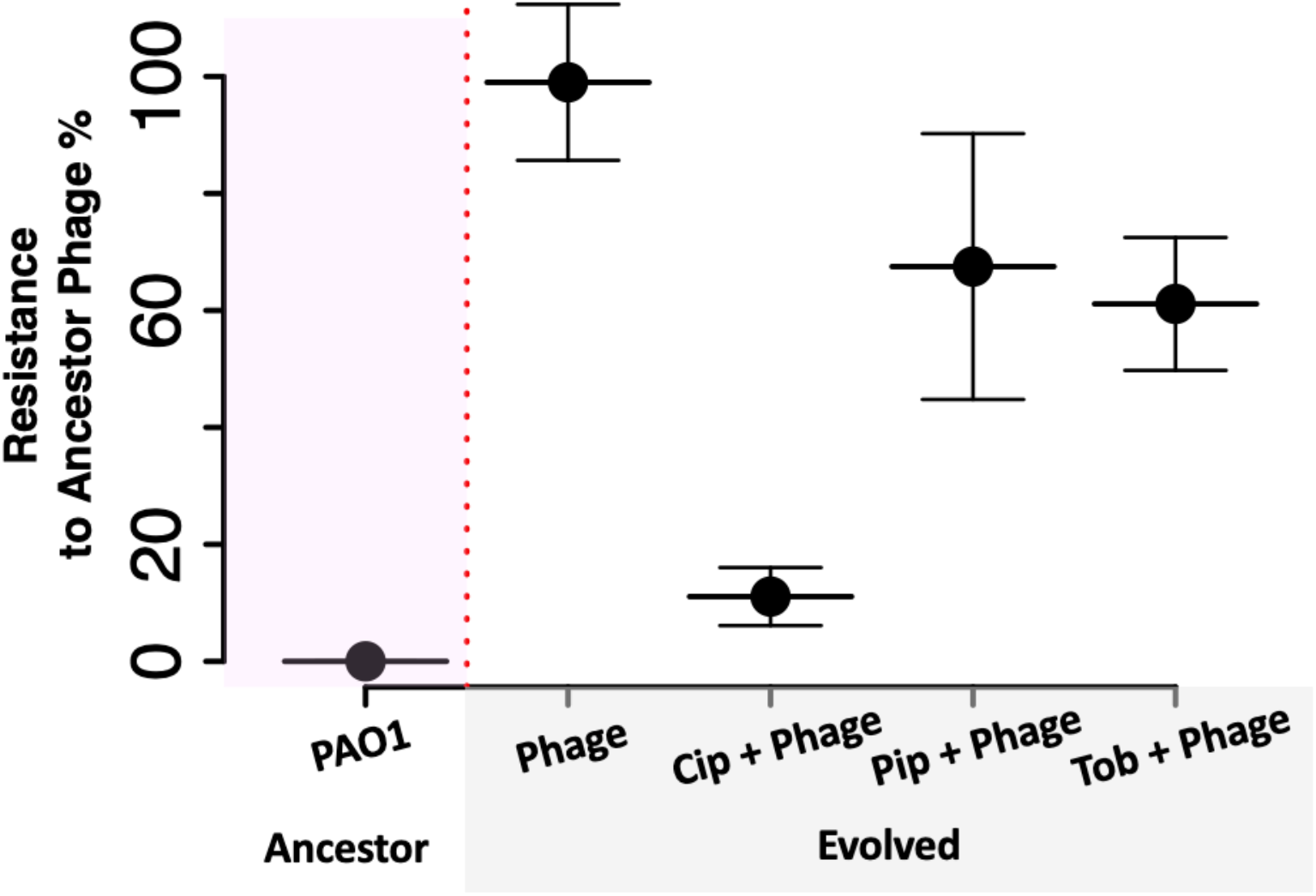
Phage combined with antibiotics promotes phage reuse. PAO1 was treated with phage OMKO1 ±3 different antibiotics for 20 days. The evolved bacteria (T10) were then examined for susceptibility to ancestral phage. Following co-evolution with phage alone, we observed high resistance to ancestral phage, consistent with ARD dynamics. With phage plus antibiotics, we observed at least partial susceptibility to the ancestral phage, consistent with FSD. The ciprofloxacin + OMKO1 line had the largest effect, approaching the initial phage impact on the ancestor (pink region). Error bars are 95% Confidence Intervals.

### Phage maintained antibiotic sensitivity during co-evolution equivalent to a clinical treatment window

Figure 4 demonstrates that the combination of OMKO1 and ciprofloxacin promotes reusability. However, OMKO1 has other already identified evolutionary advantages via its ability to reduce antibiotic resistance via antibiotic resistance counter-selection [6, 35, 36], so it is important to assess whether these advantages are compromised in this co-evolutionary context.

To assess impacts on antibiotic resistance, we determined the minimal inhibitory concentrations (MICs) of each of the replicate lines (n = 5) from each of the treatments after 20 days of co-evolution (passage T10). In the phage alone treatment, we also determined the MIC for each of the antibiotics used in the other treatments (Figure 5). As expected from OMKO1’s established ability to counter-select antibiotic resistance due to its efflux targeting activity [35], we found significantly reduced MICs across multiple phage treatments, with or without antibiotics (see SI table 3 for full ANOVA; TukeyHSD analysis). These data show the phage are still selecting against antibiotic resistance throughout the 20 day experiment, despite ongoing antibiotic re-administration and distinct co-evolutionary trajectories (ARD or FSD).

**Figure 5.**
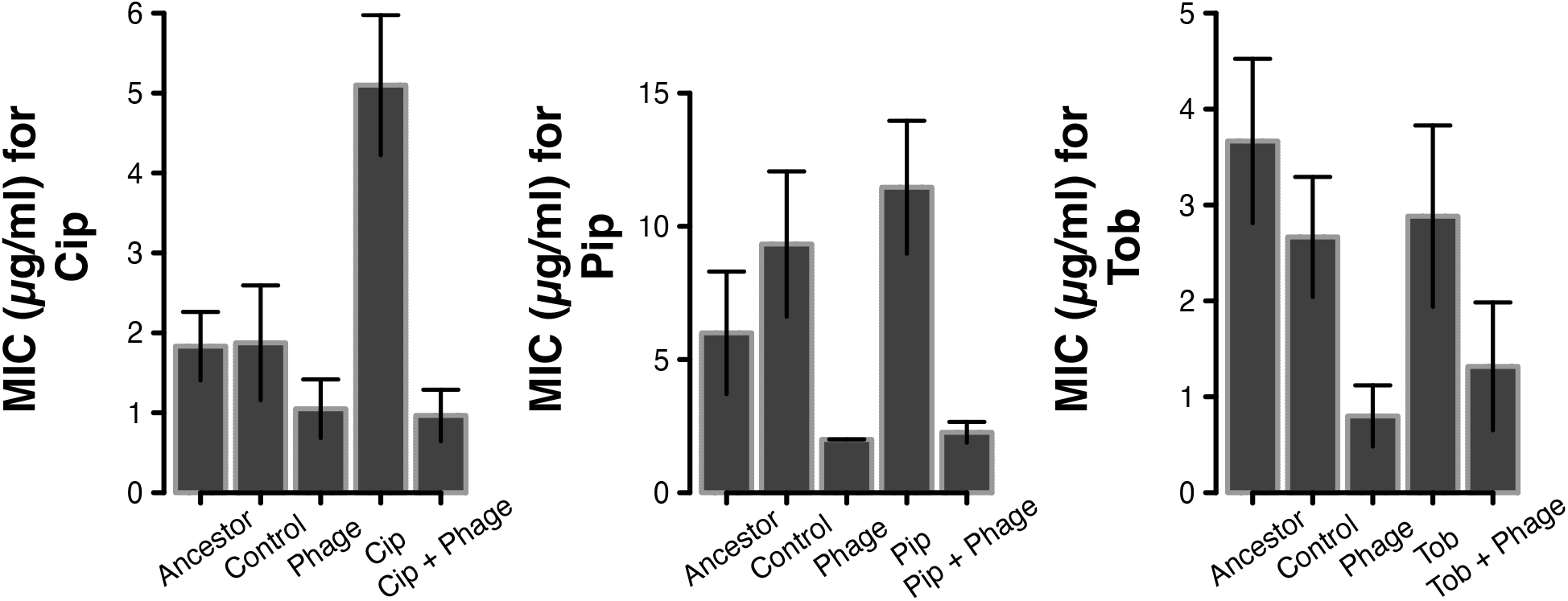
Phage maintained antibiotic sensitivity during co-evolution equivalent to a 20 day treatment window. Antibiotic sensitivity (minimal inhibitory concentration, MIC) of the ancestral bacterial strain, plus after evolution under 4 treatments: control (SCFM2 alone), phage only, antibiotic only and antibiotic plus phage. Each of the bacterial evolved lines was tested in the antibiotic treatment in which they evolved (Cip, Pip, or Tob). The Control line and the Phage line were further tested in each of the antibiotics. Error bars are 95% Confidence Intervals.

### Whole genome sequencing reveals enrichment of SNPS by treatment

Figures 2-5 demonstrate that our treatment exposures have led to significant changes in both phage and antibiotic resistances. We next turn to genomic analyses to begin to decipher the mechanistic bases of these phenotypic shifts. Given that our most promising phenotypic results involved ciprofloxacin co-administration, we focus our genomic efforts on Cip + OMKO1 evolved T10 lines plus controls. Our primary genomic hypotheses are that (1) OMKO1 will select for efflux pump modifications, to limit phage binding (at the cost of increased antibiotic susceptibility); and (2) ciprofloxacin will select for target modifications (mutations in gyrA for example [39]), efflux, or changes in envelope permeability.

The results of our whole genome sequencing analysis are summarized in Figure 6, and further detailed in the SI (SI1 Genomic analysis). In line with our expectations, following OMKO1 exposure, we found multiple efflux SNPs, and under Cip exposure we found efflux and gyrase SNPs. Surprisingly, we also found gyrase SNPs under OMKO1 exposure alone, indicating a potential role for gyrase in OMKO1 host exploitation. This is consistent with prior reports of phage requiring host gyrase to efficiently replicate their genomes [40]. In the discussion we assess the implications for potential pleiotropic constraints on PA evolution against OMKO1 plus Cip, mediated by shared efflux and gyrase targets.

**Figure 6.**
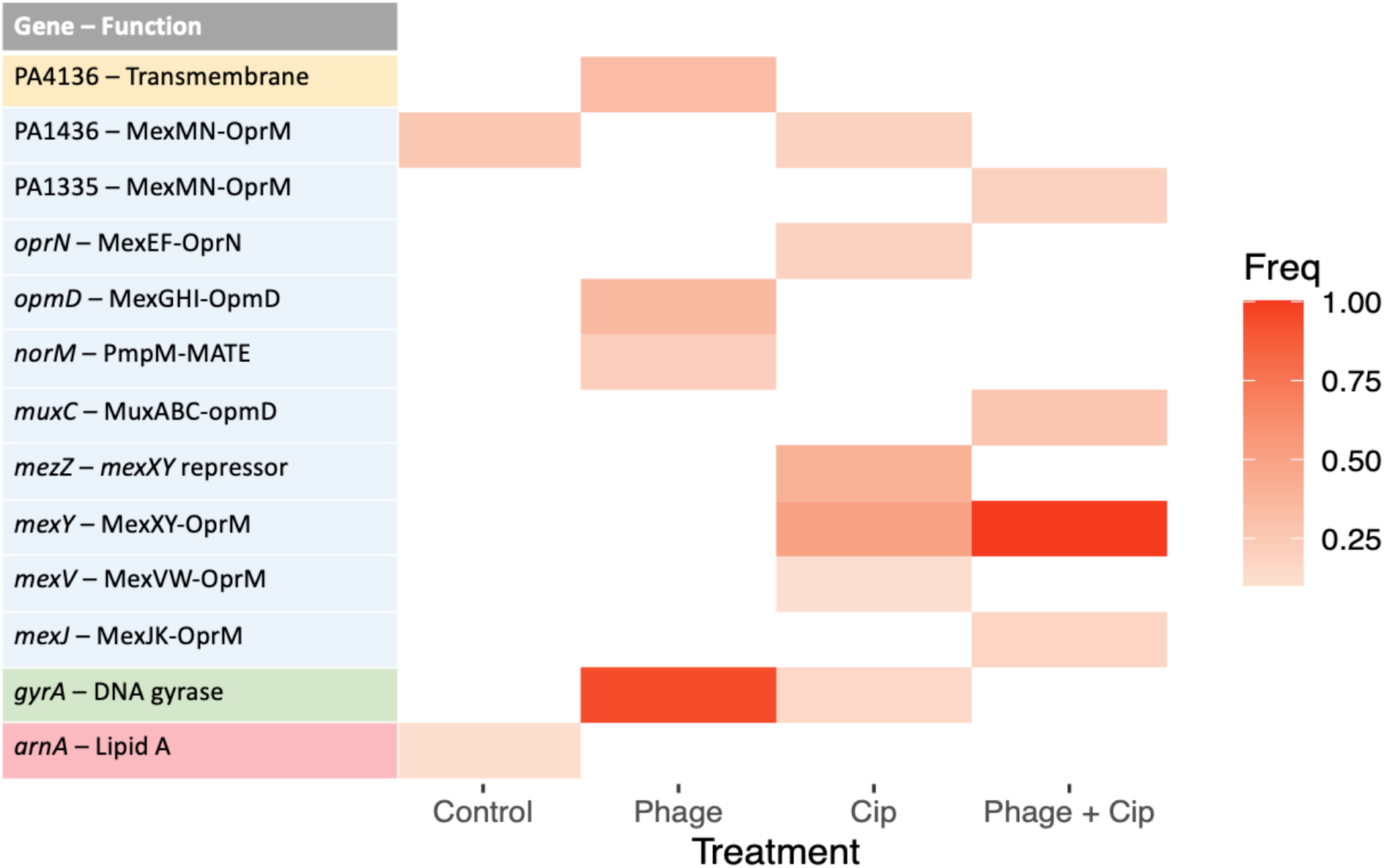
Phage OMKO1 and ciprofloxacin exposure resulted in SNPs in both efflux pumps and gyrase, independently and during co-administration. Summary of SNPs in antibiotic resistance genes in T10 isolates (versus ancestor) from the Cip arm of the co-evolution experiment (Figures 4 and 5). The table summarizes SNP frequency (cutoff > 0.1) in a curated antibiotic resistance database (CARD [41], plus ciprofloxacin target gyrase), identified by Breseq [42] from whole genome data. Genes are color coded by function: yellow is a transmembrane resistance protein, blue are efflux pumps, green is the antibiotic target (gyrase), red is membrane associated factors. (For complete sequence information, see SI1 Genomic analysis)

## Discussion

Using an experimental co-evolution approach, we demonstrate that antibiotic exposure can trigger a switch in co-evolutionary dynamics, moving from escalating arms-race dynamics, (ARD, in the absence of antibiotics) to fluctuating selection dynamics (FSD) in their presence. We pursue the clinical relevance of this finding by conducting our experiments using clinically relevant pathogens, antibiotics and phage and using a heavily benchmarked synthetic sputum medium that mirrors the conditions in chronic cystic fibrosis lung infections [32, 33]. In a chronic infection context, a major challenge is that pathogen evolution can preclude the reuse of the same therapy at future time points, narrowing treatment options with time. We reasoned that the antibiotic-induced switch to FSD co-evolution opens a window to reuse of an ancestral phage. We directly tested this prediction, finding that antibiotic co-treatment enhanced susceptibility to ancestral phage 20 days after initial phage treatment, with the greatest improvement triggered by the bactericidal antibiotic ciprofloxacin. Phenotypic and genomic characterization of evolved isolates demonstrates that efflux-targeting phage OMKO1 exerts continued selection for antibiotic susceptibility regardless of co-evolutionary dynamic or antibiotic co-treatment.

In the general context of phage-bacteria co-evolution, our results build on prior research indicating that the nature of co-evolutionary dynamics are sensitive to environmental context [23]. While prior co-evolution research has focused on the role of nutrient limitation and mutation supply [30], there are other studies that focus on bacterial evolutionary responses to phage plus antibiotic co-administration. Moulton-Brown and Friman demonstrated rapid bacterial resistance evolution to both antibiotic and phage stressors (alone and in combination), and little evidence of co-evolution, in short – reuse could fail as combined resistance occurred [43]. In contrast, we establish the importance of antibiotic stressors as clear determinants of a transition from ARD (Figure 2) to FSD (Figure 3) in a clinically relevant context, opening a principled path to overcoming the ARD barrier to phage therapy reuse.

Our choice of synthetic sputum medium (SCFM2, [34, 44]) allows us to connect controlled co-evolutionary experiments in a defined medium to a clear clinical context of chronic lung infections in people with cystic fibrosis. SCFM2 recapitulates the biochemical composition of sputum and its physical viscosity [34], and allows *P. aeruginosa* to replicate patterns of gene expression observed *in vivo* better than all other alternate models tested, including mice [33]. In this context, our results support the expectation that *P. aeruginosa* will display ARD co-evolution if treated with phage OMKO1 alone *in vivo*, presenting a barrier to reuse. If this barrier is observed in the clinic, our results also offer a simple solution – administration of OMKO1 in tandem with antibiotics will promote a transition to FSD co-evolution, therefore increasing the expected success of the same reference OMKO1 strain on secondary reuse.

We of course recognize that SCFM2 cannot replicate all elements of the *in vivo* environment, and it is possible that additional *in vivo* stressors, for example the microbiome or host immune responses, could also promote transitions to FSD co-evolution. In partial support of this possibility, the addition of protists in a phage-bacteria co-evolution study reduced but did not eliminate arms race dynamics and induced a similar lack of generalized resistance in evolved isolates [45]. Whether or not immune effectors and/or interacting microbial species can trigger a switch to FSD remains to be determined. We note at this point that our key result remains relevant despite this uncertainty – adding antibiotics can increase confidence that FSD is achieved and therefore increase the likelihood of future successful reuse.

We also recognize that other barriers to reuse exist, including the potential for host immune responses to phage treatments. Phages can trigger both innate and adaptive immune responses, which are now the subject of increased scrutiny [46], leading to a number of solutions and avenues to promote efficacy in an immunological context, such as selection for less immunogenic phage via *in vivo* passaging [47-49]. Until we know more about co-evolutionary dynamics in *vivo* we note that our strategy of antibiotic co-administration is a relatively safe and, allows for non-inferiority treatment to be given which will improve the likelihood of phage therapy application approval [50].

Our results indicate that among the three drugs tested (all commonly used in CF, [51]), ciprofloxacin most strongly enhanced treatment efficacy on reuse (Figure 4). To address the mechanism of ciprofloxacin-induced phage re-usability, we turned to whole-genome analysis (Figure 6), which revealed expected changes in efflux (the phage target) under phage treatment and gyrase (the antibiotic target) under antibiotic treatment. Previous studies demonstrate that OMKO1 requires efflux pumps to infect and selection with the phage selects for sensitive bacteria to common antibiotic treatments, even when the antibiotic is co-administered with the phage [6, 35, 36]. Our phenotypic results support this common pattern (Figure 5), as does our genomic characterization in the case of ciprofloxacin, where we see multiple efflux mutations in the presence of the efflux-binding phage (Figure 6). We also see efflux modifications in the absence of OMKO1, highlighting the key importance of efflux in governing phenotypic tradeoffs between antibiotic and phage resistances.

Mimicking the dual importance of efflux for both treatment components, our gyrase results also suggest a dual role. Both treatment with ciprofloxacin alone and phage alone both lead to SNPs within *gyrA*. GyrA is the target of ciprofloxacin and *gyrA* mutations can confer ciprofloxacin resistance [39]. In parallel, host gyrase also impacts phage replication [40, 52]. These results imply that *gyrA* presents a second pleiotropic mechanism of antibiotic versus phage resistance (in addition to efflux pleiotropy [35]) that is specific to ciprofloxacin, and potentially accounts for the more constrained evolutionary trajectory during OMKO1 plus ciprofloxacin treatment.

This study focuses on building a tractable experimental model, while maximizing clinical relevance *in vitro*. Building from this platform, key next steps are to investigate the *in vivo* dynamics of phage therapy. Understanding the mechanisms and dynamics of phage resistance evolution in patients is the crucial next step towards improving the long-term re-usability of phage therapy. Temporal samples taken from clinical applications will allow a repeat of the phenotypic and genomic characterizations demonstrated in Figures 2-6, and will help to establish the mode of bacteria-phage co-evolution *in vivo*, and as a function of different antibiotic co-administrations. With sufficiently large cohorts, additional insights may also be gained into the role of variable polymicrobial community factors, or variable immune profiles. Our focus on re-usability must not detract from the primary treatment goal which is of course infection clearance. We focus here on re-useability as a pragmatic goal given the recalcitrant nature of chronic infections. We also note that sequential reuse of a repeatedly effective treatment offers a clear path towards reduced mortality, morbidity and an increased likelihood of clearance. Finally, this work gives direct clinical relevance to understanding co-evolutionary dynamics a key component of evolutionary medicine.

## Methods

### Bacteria and bacteriophage preparation

*Pseudomonas aeruginosa* (Washington PAO1) was grown at 37 °C (constant shaking at 200 rpm) in Synthetic Cystic Fibrosis Media 2 (SCFM2) media, prepared as described in [44]. For phage amplification, 10 µl of purified OMKO1 phage [35] was added to mid-log growing PAO1 (4 hours at 37 °C), then incubated overnight (18 hours) without shaking at 37 °C (Thermofisher isotemp™). Phage were extracted by using a 10% vol/vol of chloroform, the mixture vortexed and centrifuged (1 min 10000 RCF) to separate the phases. The supernatant containing phage were pipetted into a sterile 2 ml Eppendorf and stored at 4 °C. For the complete phage preparation protocol, see [53].

### Co-evolutionary experiment

We tested the short term (10 transfers over 20 days) bacterial evolutionary responses to antibiotics and phage; 20 days was selected to match current clinical applications of phage. Bacteria were exposed to a single antibiotic (ciprofloxacin, tobramycin, piperacillin/tazobactam), in the presence of the phage OMKO1, or in the presence of both antibiotic and phage. In a control treatment, neither antibiotic nor phage were added. The experiment was conducted in 2-mL microcosms of SCMF2 in 24 well microtiter plates, incubated at 37 °C with shaking at 200 rpm (New Brunswick I 26). For each of the four treatments, 5 replicate lines were initiated in separate microcosms, for a total of 40 lines across all treatments (5 control + 5 phage + 3 x 5 Antibiotic-only + 3 x 5 phage plus antibiotic). Each microcosm was inoculated with 20 μL of PAO1 (*c*. 5 × 10^6^ cells) from an overnight population initiated from a single ancestral *P. aeruginosa* PAO1 colony. For treatments with phage, 5 μL of the ancestral phage stock (*c*. 5 × 10_6_ phage particles) were added together with the inoculated bacteria, producing a ∼1:1 ratio of bacteria and phage. For treatments with antibiotics, we used clinically reported levels of antibiotics in sputum (1 µg/ul ciprofloxacin, 2 µg/ul tobramycin, 2.5/0.3125 µg/ul piperacillin/tazobactam) [54-56]. For each microcosm, a 20 μL sample of culture (containing both bacteria and phage) was transferred to a new microcosm with fresh SCFM2 medium every 48 hours, for a total of 10 transfers (*c*. 70 bacterial generations for controls). At each transfer, a 50 μL sample was stored in 50% glycerol at −80 °C for subsequent analysis.

### Minimum inhibitory concentration of clinical antibiotics

To assess the level of antibiotic resistance of the phage-resistant bacteria, the minimum inhibitory concentration (MIC) [57] was determined for three antibiotics with different modes of action (ciprofloxacin, tobramycin, piperacillin/tazobactam). At the end of the 10 transfers, we determined the antibiotic resistance of the evolved bacteria (MIC) for 3 arbitrarily chosen colonies from each replicate line. Briefly, phage-resistant bacteria were re-streaked onto LB agar plates overnight, then grown in LB overnight to ensure phage-free cultures. Optical densities of these cultures were measured at a wavelength of 600 nm (OD_600_) and then re-suspended to a uniform OD_600_ of 1.0. In the same way, we prepared ancestral (phage-susceptible) bacterial cultures. In 96-well microtiter plates, a 2-fold dilution series of each antibiotic was prepared, starting at 8 µg/ml and finishing at 0.125 µg/ml for ciprofloxacin, 10 µg/ml to 0.156 µg/ml for tobramycin, and 25/3.125 µg/ml to 0.4/0.05 µg/ml for piperacillin/tazobactam. Bacteria were added to each well to a final OD_600_ of 0.05 and incubated at 37 °C for 18 hours. Bacteria were considered as having grown if growth was visible after 18 hours. The MIC was determined as the lowest antibiotic concentration at which no bacterial growth occurred. Three individual colonies were isolated from each replicate culture and three from the ancestor, and their MIC determined for three replicates.

### Identification of phage infectivity

For the time shift assays and reuse-phage infectivity assay a cross-streak experiment was used to determine infectivity and resistance. Bacterial isolates from each of the replicate lines and timepoints where plated on LB agar overnight. Single colonies (12) from these plates were crossed against 3 liquid lines of extracted phage for each phage time tested. In the time-shift experiment each bacterial time was streaked against all the phage times. I.e., T1 bacteria were crossed against T1, T5 and T9 phage. T5 bacteria against T1, T5 and T9 phage. As well as T9 bacteria against T1, T5 and T9 phage. This allowed a full interaction model to be tested. An isolate was considered sensitive if there was a clear interruption in the growth at each of the phage lines and scored 0. It was considered resistant if it clearly grew and scored 1. Intermediate results, partial growth in 1 or more crosses, were scored as 0.5. See method in [20] for full description.

### Statistical analysis

For the diagnosis of co-evolutionary dynamics, a generalized linear mixed effect method (GLMM) with phage time and bacterial time as fixed effects and the replicate lines as random effects was used, see [38] for detailed explanation of method. For the MIC data of bacterial resistance to the three antibiotics an Analysis of Variance (ANOVA) (R 4.03 base package, supplemented with the ‘car’ and ‘lsmeans’ packages using the Tukey adjustment) was employed.

### Whole genome sequencing and analysis

Whole genome sequencing was performed by growing evolved population in SCFM2 over night at 37 °C and extracting and purifying genomic DNA using the Promega wizard Genomic DNA purification Kit following the manufactures protocols. DNA was sequenced via Microbial genome sequencing center Pittsburgh using their in-house pipelines. SNPs were called using BRESEQ [42] and mapped against an ancestral genome deposited W-PAO1 sequence online at pseudomonas.com [58]. Using a custom R pipeline SNPs were compared across treatments and a frequency of greater than 10% was used to filter out low quantity SNPs.

## Acknowledgements

This work was funded by the NIH via both 1R21AI143296 and 1R21AI156817, the Cystic fibrosis foundation to SPB via BROWN19P0 and a Cystic Fibrosis Foundation for a Fellowship to JG GURNEY20F0. Finally, this work was also supported by Army Research Office grant W911NF-19-1-0384. We thank Michael Hochberg, Joshua Weitz, and members of the Brown lab especially John Varga for helpful discussions.

