## Supplementary material for "Building reusable phage and antibiotic treatments via exploitation of bacteria-phage coevolutionary dynamics": SI table

**SI1. statistical analysis of co-evolutionary dynamics.**

In order to diagnose the form of co-evolution between bacteria and phage, we analyzed generalized linear mixed models (GLMMs, [1]) of bacteria – phage interaction outcomes, structured by sampling time, with separate models for each of the four antibiotic treatments (Cip, Pip, Tob, or no antibiotic). Specifically, we modeled phage infectivity (‘impact’) as a function of the passaging timepoint of the phage (‘phageT’) and the passaging timepoint of the bacteria (‘bacT’), with an additional random factor to model variation across replicate lines: i.e. $Impact \sim phageT\times bacT+ randomfactor:line$. Note that we analyzed all comparisons between T1/T5/T9 bacteria and T1/T5/T9 phage, presenting a larger dataset than the data shown in Figures 2 and 3, see table S1 for model coefficients. To assess significance of the model terms, we next performed an ANOVA on each model (table S2 the ANOVA results for the phage alone treatment). The package “lme” in R does not compute P values therefore to compare outputs we performed an ANOVA on the constructed models [2].

Table S1. Co-evolutionary dynamics of OMKO1 in SCFM2. GLMM statistical analysis of time-shift experiment. Model is $Impact \sim phageT\times bacT randomfactor:line$	.

| **Term** | **numDF** | **denDF** | **F-value** | **p-value** | **Treatment** |
| --- | --- | --- | --- | --- | --- |
| (Intercept) | 1 | 32 | 34.47008 | <.0001 | phage |
| phageT | 2 | 32 | 28.96492 | <.0001 | phage |
| bacT | 2 | 32 | 6.8856 | 0.0033 | phage |
| phageT:bacT | 4 | 32 | 6.76879 | 0.0005 | phage |

Table S2. Co-evolutionary dynamics of OMKO1 in SCFM2 with different antibiotics. GLMM statistical analysis time-shift experiment. Model is $Impact \sim phageT\times bacT + randomfactor:line$

| **Term** | **numDF** | | **denDF** | **F-value** | **p-value** | **Treatment** |
| --- | --- | --- | --- | --- | --- | --- |
| (Intercept) | | 1 | 32 | 193.17039 | <.0001 | Phage + Cip |
| bacT | | 2 | 32 | 0.00245 | 0.9976 | Phage + Cip |
| phageT | | 2 | 32 | 1.24603 | 0.3012 | Phage + Cip |
| bacT:phageT | | 4 | 32 | 6.97666 | 0.0004 | Phage + Cip |
| (Intercept) | | 1 | 32 | 78.27201 | <.0001 | Phage + Tob |
| bacT | | 2 | 32 | 2.99549 | 0.0642 | Phage + Tob |
| phageT | | 2 | 32 | 1.88299 | 0.1686 | Phage + Tob |
| bacT:phageT | | 4 | 32 | 6.2428 | 0.0008 | Phage + Tob |
| (Intercept) | | 1 | 32 | 134.27051 | <.0001 | Phage + Pip |
| bacT | | 2 | 32 | 3.6015 | 0.0388 | Phage + Pip |
| phageT | | 2 | 32 | 4.1713 | 0.0246 | Phage + Pip |
| bacT:phageT | | 4 | 32 | 3.74651 | 0.0131 | Phage + Pip |

**SI2. Minimum inhibitory concentration** **analysis of T10 evolved lines.**

We preformed Minimum Inhibitory Concentration (MIC) tests on the evolved T10 bacteria (Figure 5). (See main text methods for details). These data show that a number of the treatments reduced antibiotic resistance, consistent with the known counter-selective impacts of phage OMKO1 on antibiotic efflux [3].

Table S3. Tukey analysis of ANOVA on MIC data, pre and post 20 days (T10) of coevolution (T10). Anc = Ancestor (T0), AB_= antibiotic alone lines (T10), Phage = Phage alone line (T10), PhageAB = Phage and Antibiotic (T10), Control = The evolution control line (T10, SCFM2 only - no phage, no antibiotic).

| **Contrast** | **Estimate** | **SE** | **df** | **t.ratio** | **p.value** | **treatment** |
| --- | --- | --- | --- | --- | --- | --- |
| Ab_ - Anc | 3.26666667 | 0.49489008 | 58 | 6.60079242 | 1.3509E-07 | Cip |
| Ab_ - Phage | 4.05 | 0.37410173 | 58 | 10.8259322 | 1.50E-11 | Cip |
| Ab_ - PhageAB | 4.13333333 | 0.37410173 | 58 | 11.048688 | 1.49E-11 | Cip |
| Ab_ - Control | 3.225 | 0.39679481 | 58 | 8.12762648 | 3.87E-10 | Cip |
| Anc - Phage | 0.78333333 | 0.49489008 | 58 | 1.58284308 | 0.51422485 | Cip |
| Anc - PhageAB | 0.86666667 | 0.49489008 | 58 | 1.75123064 | 0.41158284 | Cip |
| Anc - Control | -0.0416667 | 0.5122599 | 58 | -0.0813389 | 0.99998977 | Cip |
| Phage - PhageAB | 0.08333333 | 0.37410173 | 58 | 0.22275581 | 0.99943701 | Cip |
| Phage - Control | -0.825 | 0.39679481 | 58 | -2.0791603 | 0.24298801 | Cip |
| PhageAB - SCFM2 | -0.9083333 | 0.39679481 | 58 | -2.2891765 | 0.16316029 | Cip |
| Ab_con - Anc | 5.46666667 | 1.44264551 | 58 | 3.7893347 | 0.00321078 | Pip |
| Ab_con - Phage | 9.46666667 | 1.0905375 | 58 | 8.68073468 | 5.95E-11 | Pip |
| Ab_con - PhageAB | 9.2 | 1.0905375 | 58 | 8.43620695 | 1.29E-10 | Pip |
| Ab_con - Control | 2.13333333 | 1.15668969 | 58 | 1.8443437 | 0.35876113 | Pip |
| Anc - Phage | 4 | 1.44264551 | 58 | 2.77268392 | 0.05554145 | Pip |
| Anc - PhageAB | 3.73333333 | 1.44264551 | 58 | 2.58783833 | 0.0859618 | Pip |
| Anc - Control | -3.3333333 | 1.49327997 | 58 | -2.2322226 | 0.18258124 | Pip |
| Phage - PhageAB | -0.2666667 | 1.0905375 | 58 | -0.2445277 | 0.9991866 | Pip |
| Phage - Control | -7.3333333 | 1.15668969 | 58 | -6.3399315 | 3.6698E-07 | Pip |
| PhageAB - Control | -7.0666667 | 1.15668969 | 58 | -6.1093885 | 8.8308E-07 | Pip |
| Ab_con - Anc | -0.7833333 | 0.56665821 | 58 | -1.3823736 | 0.64123332 | Tob |
| Ab_con - Phage | 2.08333333 | 0.42835335 | 58 | 4.86358598 | 8.763E-05 | Tob |
| Ab_con - PhageAB | 1.56666667 | 0.42835335 | 58 | 3.65741666 | 0.00482092 | Tob |
| Ab_con - Control | 0.21666667 | 0.45433733 | 58 | 0.47688501 | 0.98916489 | Tob |
| Anc - Phage | 2.86666667 | 0.56665821 | 58 | 5.05889898 | 4.3589E-05 | Tob |
| Anc - PhageAB | 2.35 | 0.56665821 | 58 | 4.14712068 | 0.00102022 | Tob |
| Anc - Control | 1 | 0.58654698 | 58 | 1.70489328 | 0.43905195 | Tob |
| Phage - PhageAB | -0.5166667 | 0.42835335 | 58 | -1.2061693 | 0.74781461 | Tob |
| Phage - Control | -1.8666667 | 0.45433733 | 58 | -4.1085478 | 0.00115778 | Tob |
| PhageAB - Control | -1.35 | 0.45433733 | 58 | -2.9713605 | 0.0336637 | Tob |

**SI3. Genomic analysis of T10 evolved isolates.**

To assess genomic basis of phenotypic changes under OMKO1 and/or ciprofloxacin treatments, we performed whole genome sequencing on 36 populations and the ancestor (PAO1). Using 3 replicates from the T10 time point taken from the following treatment lines; the phage alone, ciprofloxacin alone, SCFM2 alone, and phage and ciprofloxacin combined treatments we extracted DNA and performed whole genome sequencing (see methods main text). Genomes were examined using Breseq [4] (mapped against the reference PAO1) and a custom pipeline in R was used to identify unique SNPs as compared to the Breseq output of the ancestor [4]. We identified antimicrobial genes in the T10 isolates using an augmented antibiotic resistance database (the CARD database [5], plus the know ciprofloxacin target *gyrA* [6]).

Here we elaborate on the SNPs presented in the main text Figure 6. We see SNPs in efflux systems in all three treatment conditions: the phage, ciprofloxacin alone, and the phage and ciprofloxacin in combination. The control lines had SNPs in identified AMR genes related to lipid modification (*arnA*) and an as yet unidentified gene PA1436 (a probable efflux transporter related to *mexMN*). In the phage alone treatment, a high frequency SNP was seen in *opmD* which forms the *mexGHI*-*opmD* efflux system [7]. Further the phage alone also selected for a SNP in *norM* (*PmpM*), an efflux pump known to confer resistance to fluoroquinolones [8]. Unexpectedly the phage also appears to have selected for a mutation in *gyrA*. Ciprofloxacin treatment led to a SNP in the target *gyrA*. We also see several SNPs in efflux pumps under ciprofloxaxin treatment (*oprN*, *mexY*, *mexV*, *mexZ*, PA1436); *mexZ* is the repressor of the *mexXY* system, loss of function here could increase resistance to ciprofloxacin. The combination of phage and antibiotic led to SNPs in 4 efflux systems and of note a SNP at 100% frequency in the *mexY* gene, which forms the *mexXY* *oprM* structure – the putative attachment target structure of the phage [3]. In the main text Figure 6 we used a cutoff value of 0.1, to filter out low frequency SNPs. Reducing the cutoff to 0 increased the number of SNP detected however the pattern of the data remains consistent (Figure S1).

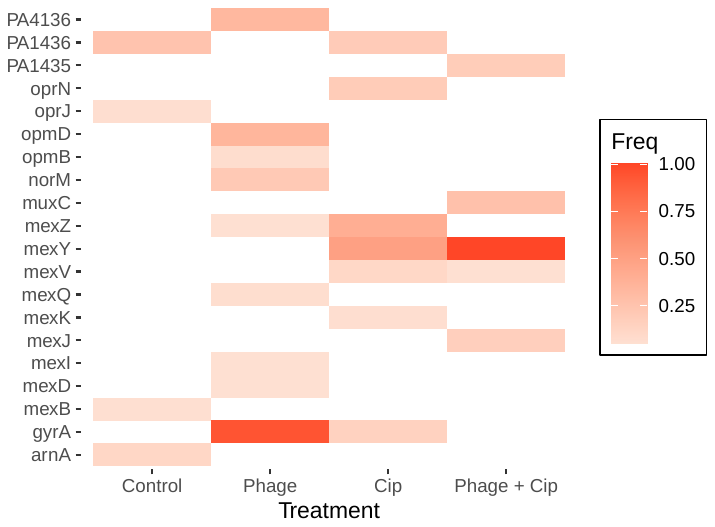

SI Figure1 **Phage OMKO1 and ciprofloxacin exposure resulted in SNPs in both efflux pumps and gyrase, independently and during co-administration (repeat of Figure 6 with frequency cutoff = 0).** Summary of SNPs in antibiotic resistance genes in T10 isolates (versus ancestor) from the Cip arm of the co-evolution experiment. The table summarizes SNP frequency (no cutoff) in a curated antibiotic resistance database (CARD [5], plus ciprofloxacin target gyrase), identified by Breseq [4] from whole genome data.
